# Hippo Signaling Modulates the Inflammatory Response of Chondrocytes to Mechanical Compressive Loading

**DOI:** 10.1101/2023.06.09.544419

**Authors:** Xiaomin Cai, Christopher Warburton, Olivia F. Perez, Ying Wang, Lucy Ho, Christina Finelli, Quinn T. Ehlen, Chenzhou Wu, Carlos D. Rodriguez, Lee Kaplan, Thomas M. Best, Chun-Yuh Huang, Zhipeng Meng

**Affiliations:** Department of Molecular and Cellular Pharmacology, Miller School of Medicine, Miami, FL; Sylvester Comprehensive Cancer Center, University of Miami Miller School of Medicine, FL; USOAR Scholar Program, Medical Education, University of Miami Miller School of Medicine, Miami, FL; Department of Biomedical Engineering, University of Miami, FL; Department of Orthopedics, University of Miami, Miami, FL; UHealth Sports Medicine Institute, University of Miami, Miami, FL

**Keywords:** chondrocyte, mechanical overload, compression, Hippo pathway, LATS inhibitor, osteoarthritis, NF-κB, PKC

## Abstract

Knee osteoarthritis (KOA) is a degenerative disease resulting from mechanical overload, where direct physical impacts on chondrocytes play a crucial role in disease development by inducing inflammation and extracellular matrix degradation. However, the signaling cascades that sense these physical impacts and induce the pathogenic transcriptional programs of KOA remain to be defined, which hinders the identification of novel therapeutic approaches. Recent studies have implicated a crucial role of Hippo signaling in osteoarthritis. Since Hippo signaling senses mechanical cues, we aimed to determine its role in chondrocyte responses to mechanical overload. Here we show that mechanical loading induces the expression of inflammatory and matrix-degrading genes by activating the nuclear factor-kappaB (NFκB) pathway in a Hippo-dependent manner. Applying mechanical compressional force to 3-dimensional cultured chondrocytes activated NFκB and induced the expression of NFκB target genes for inflammation and matrix degradation (i.e., IL1β and ADAMTS4). Interestingly, deleting the Hippo pathway effector YAP or activating YAP by deleting core Hippo kinases LATS1/2 blocked the NFκB pathway activation induced by mechanical loading. Consistently, treatment with a LATS1/2 kinase inhibitor abolished the upregulation of IL1β and ADAMTS4 caused by mechanical loading. Mechanistically, mechanical loading activates Protein Kinase C (PKC), which activates NFκB p65 by phosphorylating its Serine 536. Furthermore, the mechano-activation of both PKC and NFκB p65 is blocked in LATS1/2 or YAP knockout cells, indicating that the Hippo pathway is required by this mechanoregulation. Additionally, the mechanical loading-induced phosphorylation of NFκB p65 at Ser536 is blocked by the LATS1/2 inhibitor Lats-In-1 or the PKC inhibitor AEB-071. Our study suggests that the interplay of the Hippo signaling and PKC controls NFκB-mediated inflammation and matrix degradation in response to mechanical loading. Chemical inhibitors targeting Hippo signaling or PKC can prevent the mechanoresponses of chondrocytes associated with inflammation and matrix degradation, providing a novel therapeutic strategy for KOA.

## Introduction

Knee osteoarthritis (KOA) is a prevalent degenerative joint condition affecting about 30% of individuals over the age of 60 in the United States ^1^. The exact pathophysiology of osteoarthritis (OA) is multifaceted and not yet fully understood. Risk factors, including obesity, aging, joint trauma, and female gender, have been correlated with the development and progression of KOA ^2-5^. Obesity is considered one of the greatest modifiable risk factors for OA ^6^. The increased mechanical loading on weight-bearing joints due to obesity contributes to the KOA development and progression ^6,7^. However, the mechanisms by which mechanical loading triggers KOA pathogenesis remain to be defined for developing effective therapeutic strategies ^8^.

Mechanotransduction signaling pathways of chondrocytes sense increased loading and subsequently trigger the production of cytokines and metalloproteinases, leading to the degradation of surface articular cartilage ^9-13^. The major role of articular cartilage within the joint is to provide a smooth, lubricated surface for articulation as well as to facilitate the transmission of applied forces with minimal frictional loads ^14^. Notably, articular cartilage is avascular and devoid of lymph tissue, which limits its ability to adequately repair subsequent injury ^15-18^. The knee joint experiences numerous compressive loading cycles throughout the day. Compressive forces can be static, such as when standing, or dynamic, such as with acting movement or jumping ^19^. Chondrocytes within the cartilage sense and respond to the compressive loads by regulating and secreting essential cartilaginous extracellular matrix (ECM) components, including collagens and proteoglycans ^20,21^. Mechanical overload can lead to inflammatory conditions within the tissue through chondrocyte signaling, resulting in cartilage destruction and chondrocyte apoptosis ^22^.

Understanding the mechanisms of mechanotransduction is crucial for developing new therapies that prevent cartilage destruction and chondrocyte apoptosis. The Hippo pathway, first discovered in *Drosophila*, has been recognized as a conserved signaling pathway in regulating organ development, regeneration, and carcinogenesis *via* integrating biochemical and mechanical cues that reshape cellular transcription programs ^23-26^. The core of the mammalian Hippo pathway is a kinase cascade of the mammalian sterile twenty kinases ½ (MST1/2) and the large tumor suppressor kinase ½ (LATS1/2). Environmental, biochemical, and biomechanical signals can act through this kinase cascade to phosphorylate and inactivate YAP and TAZ, two transcription co-factors that initiate the expression of genes for cell cycling, mobility, and stemness. While the Hippo pathway effector YAP has been implicated in the development of OA, its precise pathogenic roles appear to be context-dependent and require further clarification ^27,28^.

The Hippo signaling is widely recognized as the best-characterized pathway for mechanosensing and mechanotransduction ^13,29-32^. Given that OA can be induced by mechanical overload, it is essential to understand the exact functional roles of the Hippo signaling in the development of OA. Therefore, our study utilized a model of mechanical loading on 3D-cultured chondrocytes to investigate the regulation and functional significance of LATS1/2 and YAP/TAZ in the response of chondrocytes to overload. We discovered that proper Hippo signaling is required for mechanical loading to induce inflammation and matrix degradation through a Protein Kinase C (PKC)/ Nuclear factor-kappaB (**NF**κ**B**) signaling axis. We further showed that the functional interplay between PKC and Hippo signaling is required for NFκB-activated inflammation and matrix degradation and thus serves as a new target for KOA therapeutic intervention.

## Results

### Mechanical overload alters the expression of KOA-associated genes

We established both 2- (2-D) and 3-dimensional (3-D) *in vitro* models to study the gene transcriptional responses of chondrocytes to mechanical overload. We focused on the mRNA levels of two key OA genes, interleukin-1 beta (*IL1*β) and A disintegrin and metalloproteinase with thrombospondin motifs 4 (*ADAMTS4)*, associated with inflammatory responses and extracellular matrix degradation, respectively. *IL1*β is a cytokine that induces articular cartilage degradation and is elevated in OA. *ADAMTS4* is an aggrecanase that cleaves aggrecan, the major proteoglycan found in articular cartilage, which can lead to loss of cartilage function and destruction in OA ^33,34^.

For the compression study with 2-D culture, we followed a well-established protocol ^35^. We seeded C28/I2 human chondrocytes onto trans-well membranes with 0.4 μm pore and covered the chondrocytes with agarose discs as water-permeable buffering. When the cell density reaches 80%, an 8.9g stainless-steel cylinder was placed on top of the agarose disk for 4 hours (**Fig. 1A**). This level of mechanical compression has been shown to significantly impact cell-matrix interaction and cell mobility. However, this mechanical compression cannot significantly increase the expression of *IL1*β and even reduce the expression of *ADAMTS4* in these 2-D cultured chondrocytes (**Fig. 1B**).

**Figure 1.**
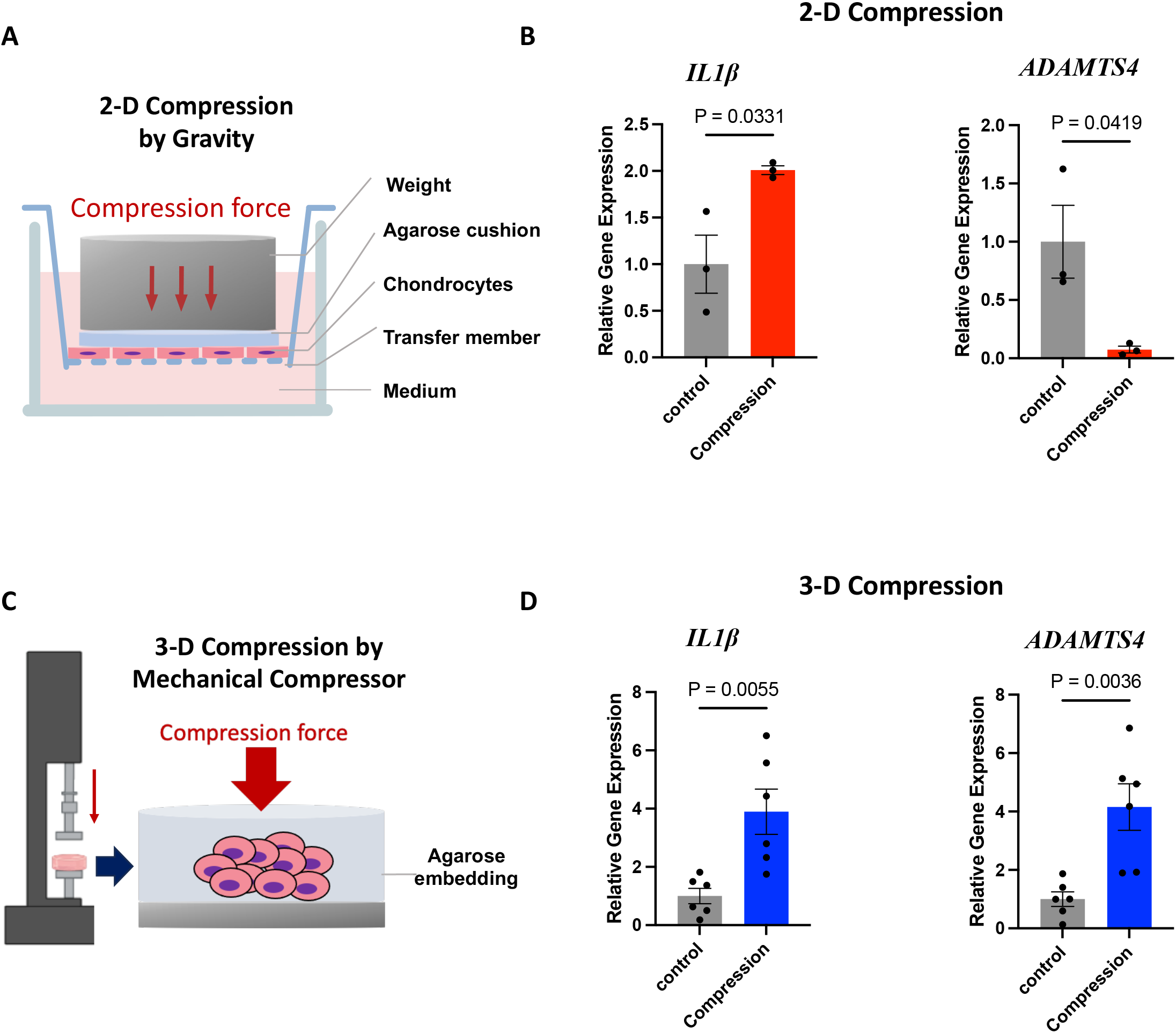
Differential responses of chondrocytes to mechanical compression in 2-dimensional (2-D) and 3-dimensional (3-D) culture. **A**. A diagram of 2-D compression generated by gravity. **B**. Quantitative Real-time PCR analyses of IL1β and ADAMTS4 gene expression in the 2-D compression model. **C**. A diagram of 3-D compression generated by a mechanical compressor. **D**. Gene expression analyses in the 3-D compression model.

For 3-D culture, we embedded chondrocytes into agarose discs and applied compression to the agarose discs using a compression bioreactor (**Fig. S1A, Fig. 1C**) to induce deformation that mimics mechanical overload caused by obesity as described previously ^36^. Twenty percent strain was used to evaluate the association between strain magnitude and cell mechano-response. With this model, we consistently observed that mechanical loading significantly induces the expression of *IL1*β and *ADAMTS4* (**Fig. 1D**), suggesting that this 3-D model well represents the pathogenesis process in vivo and can be used to define the mechanisms by which mechanical overload activates transcriptional programs for inflammation and matrix degradation.

### Mechanical overload activates NFκB and Hippo signaling without triggering cell death pathways

NFκB plays a crucial role in OA-associated processes, including synovial inflammation and chondrocyte catabolism ^1,8,37^. The NFκB family of transcription factors activates the expression of inflammatory cytokines (i.e., *IL1*β) and matrix-degrading enzymes (i.e., *ADAMTS4*) ^38,39^. It is speculated that NF-κB can be directly activated by mechanical loading ^40,41^. This notion is consistent with our QRT-PCR results of *IL1*β and *ADAMTS4* gene expression (**Fig. 1D**). We, therefore, tested whether NFκB can be activated by mechanical compression in our 3-D model. Mechanical loading strongly increases Serine 536 (S536) phosphorylation of the classical NFκB member p65 but does not affect Serine 529 phosphorylation (**Fig. 2A**). S536 phosphorylation of p65 is well known to promote the transcriptional activity of p65 independent of IκB, the endogenous negative regulator of p65 ^42-44^. However, we did not observe increased phosphorylation of IκB or increased proteolysis of p105, indicating that mechanical overload may act through a non-classical NF-κB mechanism to phosphorylate p65 at S536 and thus activate p65 ^44^.

**Figure 2.**
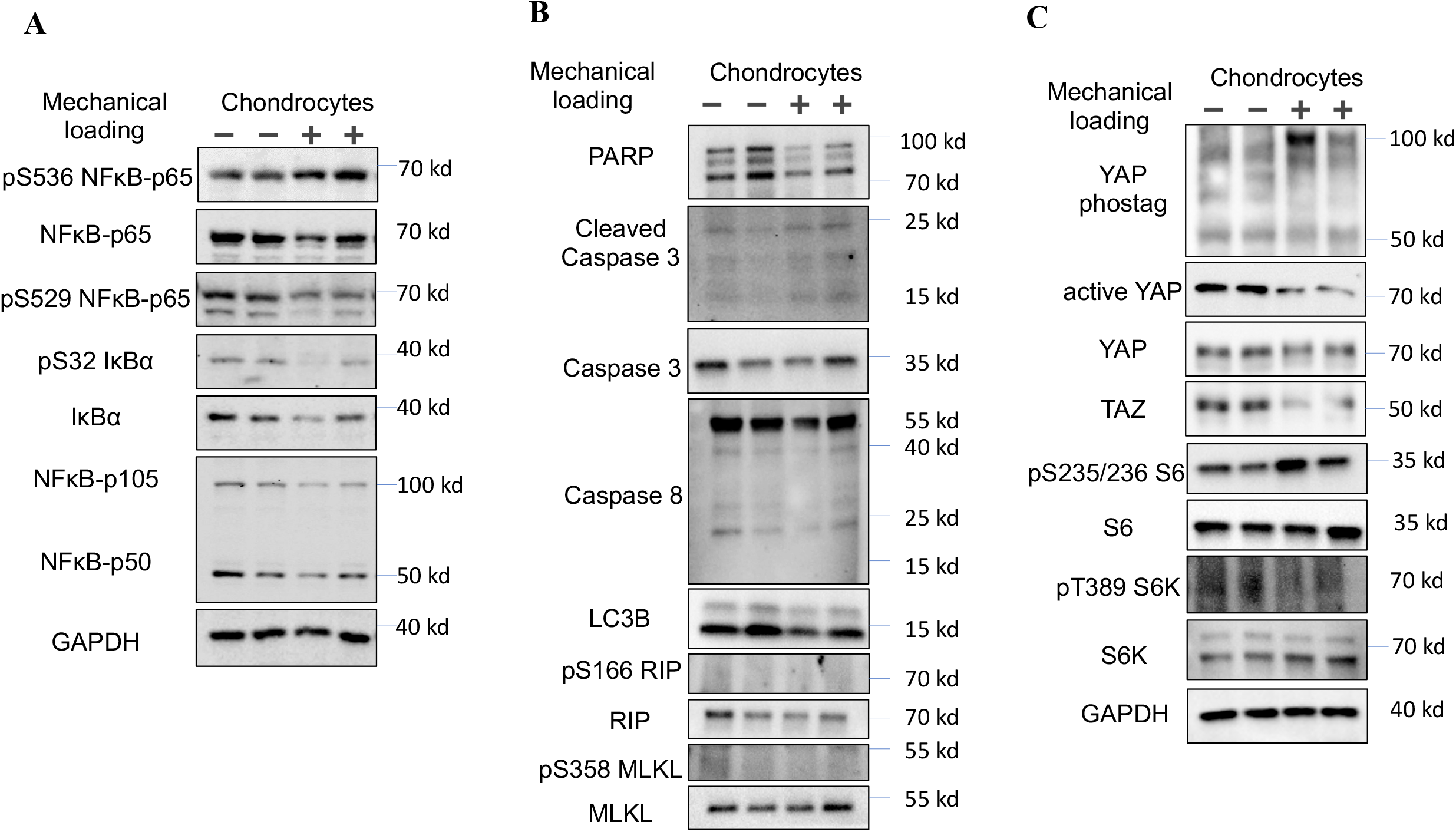
Mechanical compression activates NF-κB and Hippo signaling in chondrocytes. **A**. Mechanical compression induces phosphorylation of NF-κB p65 at Serine 536. **B**. The mechanical compression applied to chondrocytes does not lead to cell death. **C**. Mechanical compression controls YAP phosphorylation. For YAP phos-tag analysis, phosphorylated YAP migrates more slowly on this gel. Therefore, the bottom band indicates hypophosphorylated (active) YAP while the upper bands indicates hyperphosphorylated (inactive) YAP.

We also examined whether mechanical compression results in any form of cell death. We therefore first profiled chondrocytes for their apoptosis but we did not observe increased cleavage of PARP, Caspase 3, or Caspase 8, which are landmark events of apoptosis ^45^ (**Fig. 2B**). In addition, we did not observe increased levels of autophagy and necroptosis by checking cleavage of LC3B and phosphorylation of RIP and MLKL ^46,47^. These results also helped us rule out the possibility that the NFκB activation induced by mechanical loading is secondary to apoptosis, autophagy, or necroptosis caused by compression and cell deformation.

As YAP, the Hippo pathway effector is the best-characterized mechanotransduction transcription factor and is also implicated in NFκB regulation during OA, we therefore investigated YAP regulation by mechanical loading ^27,28^. The activity of YAP is mainly determined by its phosphorylation status ^13,29-32^. Therefore, we performed a Phos-tag electrophoresis where an up-shift of target protein bands indicates its hyperphosphorylation. We found that mechanical loading strongly induced YAP phosphorylation (**Fig. 2C**). Consistently, the mechanical loading decreased the active YAP, which was detected with an antibody recognizing non-phosphorylated YAP proteins, and the total protein level of TAZ, which is negatively correlated with its phosphorylation status. Besides Hippo signaling, there have been reports indicating mTORC1 signaling can also respond to mechanical load in some scenarios ^48,49^. However, we did not observe an increased mTORC1 activity, shown by the phosphorylation of mTORC1 substrate S6K and its downstream S6 at specific residues.

### Physiological biomechanical microenvironment is essential for proper regulation and activity of Hippo signaling in chondrocytes

Compared to 2-D monolayers, 3-D agarose culture models may better maintain the chondrocyte phenotype and better mimic the physiological microenvironment under compression ^50^. We, therefore, speculate that the biomechanical conditions (3-D vs. 2-D) may regulate the Hippo pathway activity in chondrocytes. In fact, our results showed significant differences in the expression of four classical YAP target genes (*CTGF, CYR61, ANKRD1*, and *AMOTL2*) between 2-D and 3-D cultures (**Fig. 3A**). All genes in the 2D culture environment were higher when compared to the 3D cultures, which were grown in a softer matrix and in an anchorage-independent status that recapitulate the features of a normal knee microenvironment ^51^. These results are consistent with the notion that cell attachment to stiff matrix activates YAP by inactivating the Hippo pathway ^52,53^. Therefore, we next determined the Hippo kinase activation status by examining the phosphorylation of LATS kinase at its phosphorylation motif (pLATS-HM), which is substantially higher in the 3D cultures than that in 2D-cultured chondrocytes (**Fig. 3B**). Consistently, phosphorylation of YAP, as shown by our phos-tag electrophoresis analyses, is also strongly increased at 3D compared with at 2-D.

**Figure 3.**
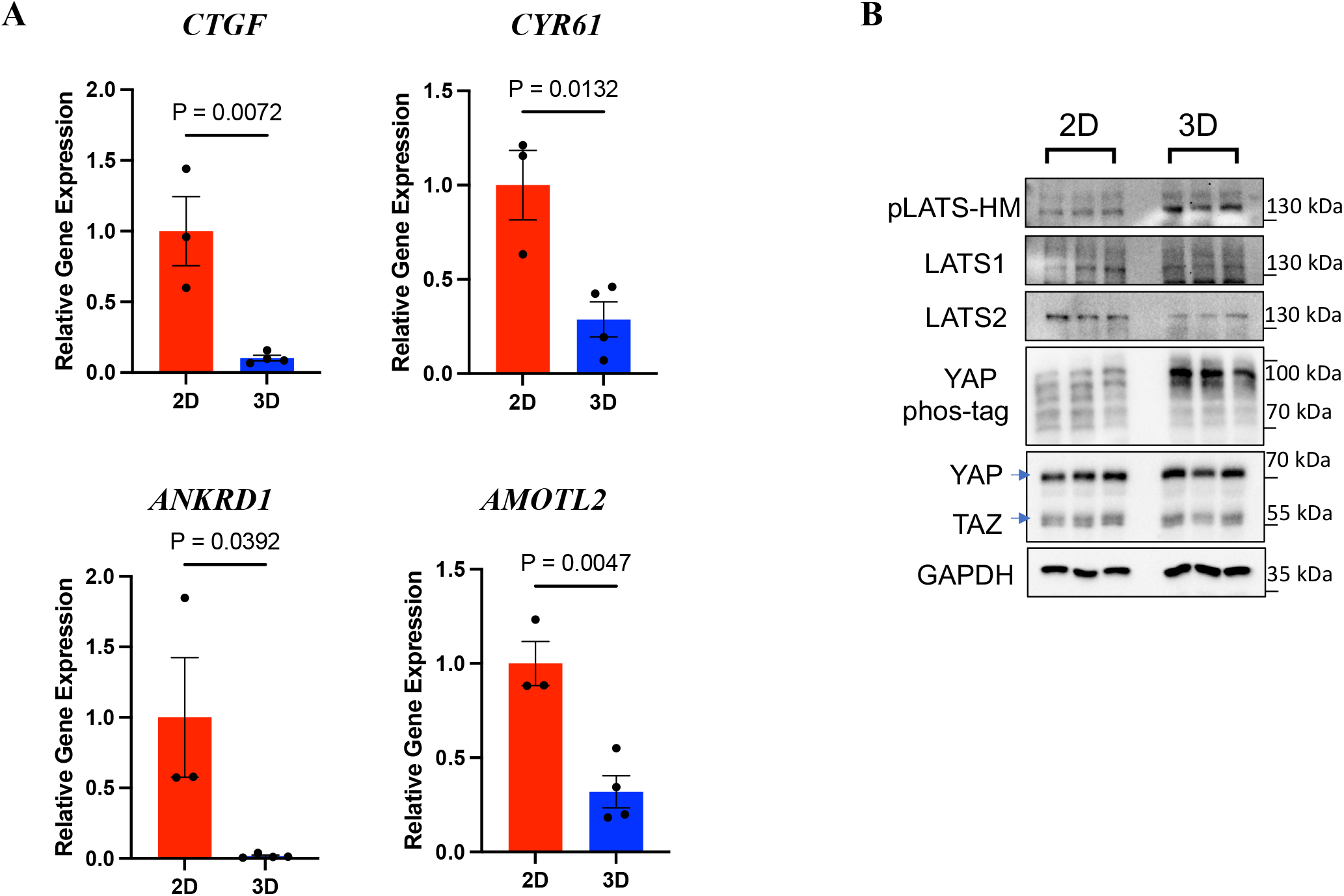
Differential activation status of the Hippo pathway in 2-dimensional (2-D) and 3-dimensional (3-D) culture. **A**. Quantitative Real-time PCR results for the relative CTGF, CYR61, ANKRD1, and AMOTL2 gene expression of chondrocytes under 2-D or 3-D culture conditions. **B**. Immunoblot for the phosphorylation of Hippo pathway core components LATS1/2 and YAP.

The activity of the Hippo pathway is normally high to maintain the quiescence of cells and prevent aberrant cell proliferation. To evaluate the Hippo pathway activity in the 2-D and 3-D chondrocyte cultures, we used CRISPR/Cas9 to delete LATS1/2 or YAP genes in chondrocytes (**Fig. 4A, B**). Lentiviruses expressing Cas9 and sgRNAs for LATS1, LATS2, or YAP were used to infect chondrocytes to generate knockout cell pools for each gene. The LATS kinase activity is tightly regulated by cell confluence ^29,54^. Our Phos-tag electrophoresis data clearly showed that deletion of both LATS1 and LATS2 (sgLATS) strongly comprised YAP protein phosphorylation induced by cell confluence (**Fig. 4A**), affirming the deficiency in the Hippo kinase cascade. Moreover, we achieved highly efficient gene knockout of YAP in the cells (**Fig. 4B**).

**Figure 4.**
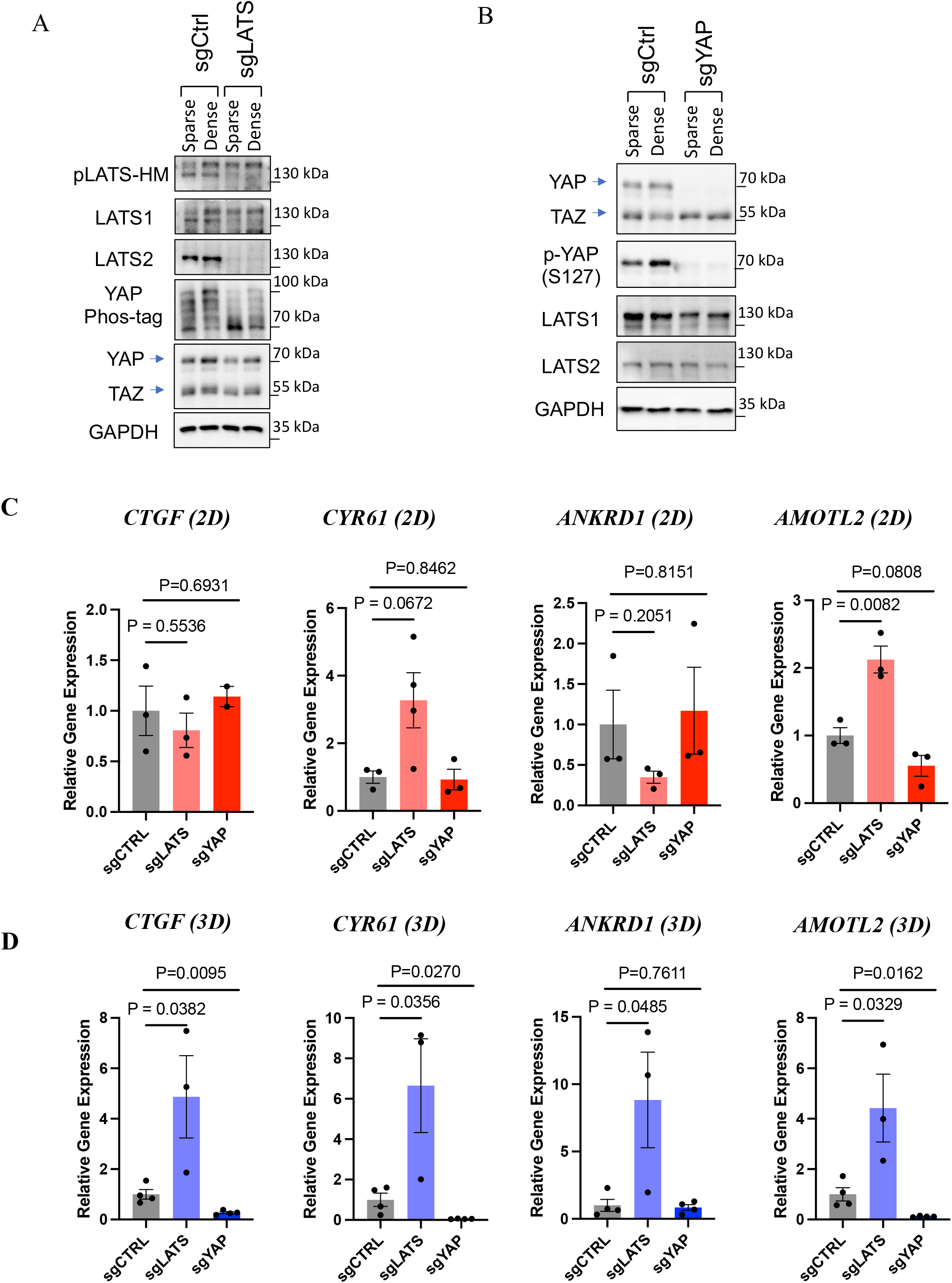
Deletion of LATS1/2 and YAP/TAZ by CRISPR/Cas9 in chondrocytes. **A**. Immunoblot showing the expression and/or phosphorylation of LATS1/2 and YAP/TAZ in chondrocyte pool with the expression of Cas9 and sgRNAs targeting LATS1/2. **B**. Expression or phosphorylation of YAP or TAZ after CRISPR/Cas9-mediated gene deletion. **C, D**. Quantitative Real-Time PCR of Yap target genes at 2D or 3D.

Surprisingly, deletion of LATS1/2 cannot further enhance the expression of YAP target genes (*CTGF, CYR61*, and *ANKRD1)*, except *AMOTL2*, at 2-D culture (**Fig. 4C**). This is likely because the activity of LATS1/2 was already very low due to the stiffness of 2-D cultured chondrocytes, which is known to inactivate LATS1/2 and thus activate YAP ^53,55^. On the other hand, deletion of YAP alone cannot reduce the expression of some YAP target genes (**Fig. 4C**), which may be due to the compensatory activation or over-expression of TAZ in the YAP knockout cell pool (**Fig. 4B**). Contrary to the 2-D culture, the chondrocytes at the 3-D culture maintain the Hippo pathway activity (**Fig. 4D**). Deletion of LATS1/2 (sgLATS) can robustly increase the expression of YAP target genes (*CTGF, CYR61, ANKRD1*, and *AMOTL2*) in 3-D cultured chondrocytes. It should be noted that, though the activity of YAP is much lower at 3-D than that at 2-D, chondrocytes still maintain a certain level of YAP activity at the 3-D culture, as deletion of YAP further reduced the expression of the YAP target genes (*CTGF, CYR61*, and *AMOTL2*).

In summary, our results suggested the artificial condition of 2D cultures may abnormally activate YAP in chondrocytes by inactivating the Hippo kinases LATS1/2 (**Fig. 3**,**4**).

### Proper regulation and activity of Hippo signaling is required for mechanoresponses of NFκB to mechanical loading

Given that Hippo signaling serves as a mechanical checkpoint and sensor of cellular homeostasis in response to biophysical environmental change, we hypothesize that deficiencies of Hippo signaling could lead to deficient mechanoresponses of chondrocytes to mechanical loading.

We first focused on whether deletion of LATS1/2 or YAP can oppositely regulate NFκB activation by mechanical loading. Unexpectedly, mechanical loading failed to induce p65 S536 phosphorylation or expression of target genes (*IL1*β and *ADAMTS4)* in LATS KO chondrocytes (sgLATS1/2) and YAP KO (sgYAP) as in the control cells (sgCTRL) (**Fig. 5A, B**).

**Figure 5.**
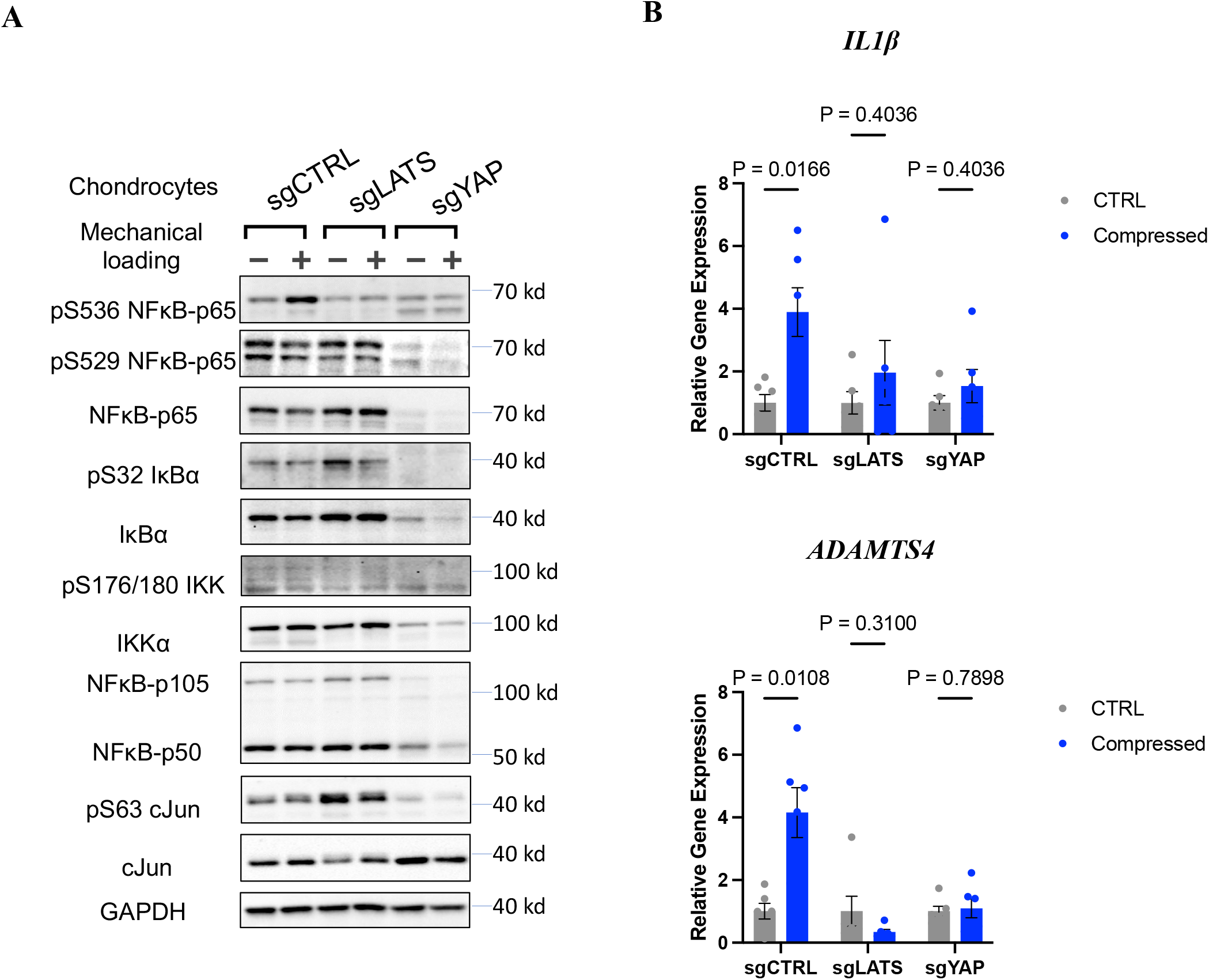
Hippo pathway is required for mechanical loading-induced NF-κB activation. **A**. Deletion of either LATS or YAP abolishes NF-κB p65 phosphorylation at Serine 536. **B**. The expression of NF-κB target genes IL1-β and ADAMTS-4 in the control, LATS1/2-KO (sgLATS1/2), and YAP-KO (sgYAP).

Detailed signaling pathway analyses of mechanically loaded chondrocytes revealed that the effects of LATS1/2 or YAP gene deletion on p65 S536 phosphorylation in chondrocytes are likely independent of canonical NFκB upstream regulators (IKKα/β, IκBα) as we did not observe phosphorylation of IKKα/β and IκBα, which indicates their activity, showed a similar pattern with p65 S536 phosphorylation across different treatment or knockout groups. Interestingly, the protein level of p65 was diminished in YAP KO chondrocytes.

AP1 family members, especially cJun, is well known to functionally interact with NFκB transcriptional activation ^56^, and Hippo signaling has recently been reported to tightly control c-Jun and other AP1 members in cellular acute response ^57^. Therefore, we also examined c-Jun protein level and phosphorylation. However, though we observed differential c-Jun phosphorylation in LATS KO and YAP KO cells, which is consistent with our previous report ^57^, mechanical loading itself does not alter c-Jun protein level or activity (**Fig. 5A**). This indicates that other NFκB upstream regulators are dysregulated in LATS1/2 KO and YAP KO cells. Nevertheless, our results support our notion that the core Hippo pathway components, such as LATS1/2 and YAP, are required for chondrocytes to respond to mechanical compression.

### LATS1/2 kinase inhibitors can be potential therapeutic agents for KOA

Based on the observation in the LATS1/2 knockout cells, we explored the potential of using LATS1/2 inhibitors to prevent the inflammatory and catabolic responses of chondrocytes to mechanical compression. We, therefore, tested a LATS-2 small molecule inhibitor (Lats-IN-1, 10 μM), which was supplemented 18 hours before exposure to a 4-hour static compressive load of 20% strain. The expression of YAP/TAZ target genes (*CTGF, CYR61, AMOTL2*, and *ANKRD1*) was significantly upregulated in the compressed discs with LATS1/2 inhibitor compared to control, indicating that Lats-IN-1 can effectively inhibit the activity of LATS1/2 (**Fig. 6A**). More importantly, Lats-IN-1 suppressed upregulation of *IL1*β and *ADAMTS4* in discs subjected to mechanical load, implying that LATS1/2 inhibition can be a potential therapeutic strategy for KOA (**Fig. 6B**), which currently lacks effective treatments.

**Figure 6.**
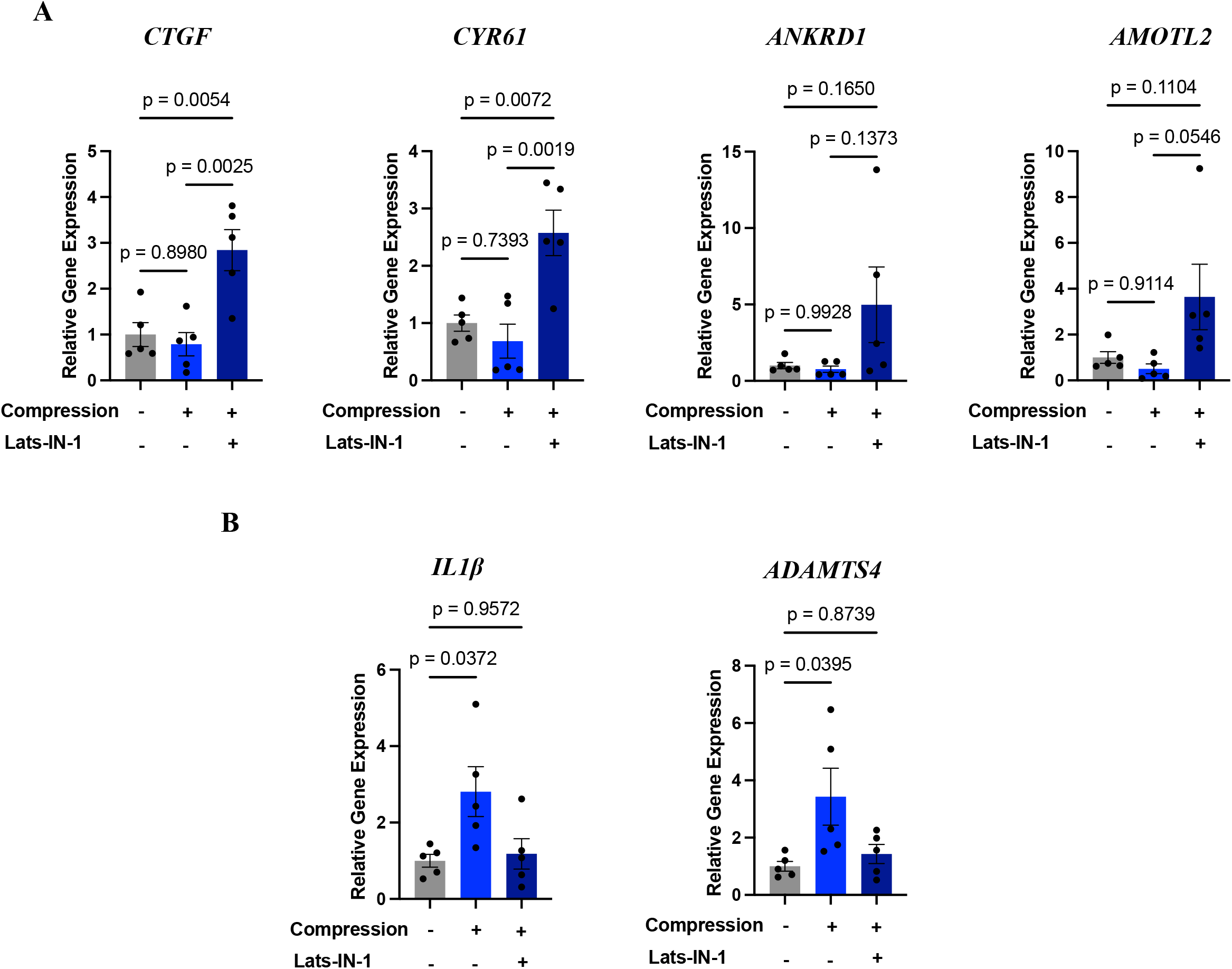
Effects of LATS1/2 inhibitors on compressed chondrocytes. **A**. The expression of YAP target genes CTGF, CYR61, ANKRD1, and AMOTL2 in the chondrocytes with LATS1/2 inhibitor and/or compression. **B**. The expression of IL1β and ADAMTS-4 in the same set of chondrocytes. *: > other two groups.

### Hippo signaling modulates the mechanoresponses of NF-κB via PKC

To understand the mechanisms by which Hippo signaling controls the responses of NFκB to mechanical loading, we profiled multiple pathways that are known to be regulated by mechanical cues or control NFκB activity (**Fig. S2**). Protein Kinase A (PKA), AKT, and AMPK are known to respond to mechanical cues, regulate chondrocyte metabolisms, and interact with NFκB ^8^. We, therefore, used their phosphor-substrate antibodies to determine whether their activities are altered in LATS KO and YAP KO cells. Mechanical compression slightly increases PKA activity in the control and LATS KO cells, though PKA activity in the latter was high even at the basal level. In the YAP KO cells, PKA activity was downregulated by mechanical loading. Therefore, the pattern was not very consistent with NFκB activity (**Fig. S2A**). Activities of AKT and AMPK do not correlate with NFκB activity well, either. Similarly, we did not observe activities of CDKs and ATM/ATR, which functionally interact with NFκB in various other settings ^56^, correlate with NFκB activity in the chondrocytes under mechanical loading (**Fig. S2B**). It should be emphasized that the effects of mechanical loading on these pathways were not very robust.

In the contrary, the activities of PKC in the control, LATS1/2 dKO, and YAP KO chondrocytes are similar to the patterns of NFκB (**Fig. 7A**). More importantly, the mechanical load can strongly stimulate PKC activities, particularly on its substrates with a molecular weight of ∼80 kd, which likely represents autophosphorylation of PKC (marked by *). PKC is well known to be a key regulator of NFκB in cardiomyocytes, which are also tightly regulated by mechanical loading. Multiple isoforms of PKCs regulate NFκB in both IKK-dependent and independent manners ^58^. In particular, atypical PKC (PKC ζ) can directly phosphorylate p65 S536 and elevate p65 transcriptional activity ^59^.

**Figure 7.**
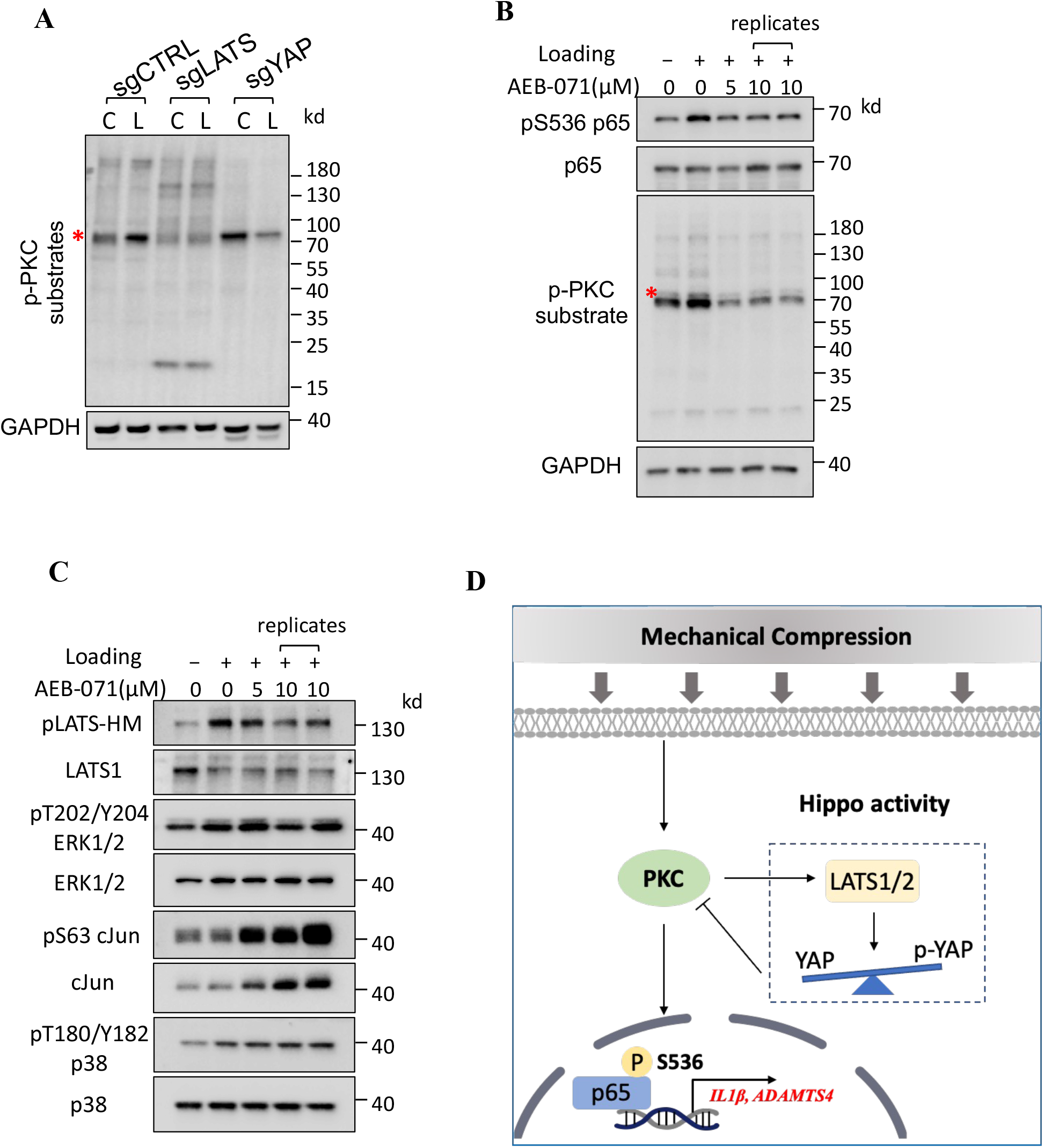
PKC is activated by mechanical compression in a Hippo-dependent manner and required for NF-κB p65 phosphorylation at Serine 536. **A**. Detection of PKC activity using phospho-PKC substrate antibodies under mechanical loading. **B**. The PKC inhibitor AEB-071 blocks NF-κB p65 S536 phosphorylation induced by mechanical compression. **C**. AEB-071 also reduces LATS activity, as shown by phosphorylation of LATS hydrophobic motif. **D**. A proposed model for the interplay of NF-κB, PKC, and Hippo signaling upon mechanical compression.

To determine the roles of PKCs in the mechanoregulation of NFκB, we applied a pan-PKC inhibitor Sotrastaurin (AEB071) to mechanically loaded chondrocytes (**Fig. 7B**). AEB071 can effectively block the phosphorylation of both PKC substrates and p65 S536, demonstrating that PKC kinase activity is required by mechanical loading to activate NFκB. Besides NFκB, Hippo, and MAPK (ERK1/2, p38, and JNK) signaling is known to be activated by mechanical loading. We further found that AEB071 also blocked LATS phosphorylation, but not increased ERK and p38 phosphorylation, resulting from mechanical loading (**Fig. 7C**).

## Discussion

### Modeling of mechanical overload-induced KOA

Several studies have been conducted to quantify *in vivo* strain levels to determine normal ranges of compression in human knee cartilage. *Eckstein et al* demonstrated a range of strain levels during typical activities, such as knee bending, jumping, and static exercise, to be within the range of 2%-10% using MRI evaluation and 3D image analysis ^19^. Previous MRI studies have shown that cartilage deformation in knees increased with a higher BMI ^60^ and is greater in osteoarthritic knees than in healthy knees ^61^. Furthermore, it was found that 25% static compression for 4 hours resulted in a transient increase in chondrocyte’s *ADAMTS4* expression and a 10% strain was modeled as a normal strain showing no effects on gene expression ^36^. We, therefore, developed our model to study mechanotransduction signaling and transcriptome using a 4-hour 20% compression loading regimen. On one hand, our model has successfully recapitulated the mechanoregulation of NFκB and subsequent target gene expression. On the other hand, we have been using conventional cell culture medium and atmospheric conditions, which are rather different from the nutrient and oxygen levels in the cartilage. As mechanosensing pathways, such as Hippo and NFκB signaling, also incorporate biochemical cues for cellular homeostasis, therefore we will further optimize our system using a human plasma-like medium and more advanced cell culture incubator with oxygen and hydrostatic pressure control ^62^.

Since dynamic loading of low magnitude can suppress *IL1*β-mediated catabolic responses of chondrocytes^63-66^, dynamic compression studies utilizing a similar protocol could also provide important findings to evaluate the involvement of the Hippo pathway under different loading conditions, which may model physiologic environments in a more realistic occurrence. Other loading factors, such as loading duration, magnitude, and frequency, should also be further defined. Furthermore, compression rate, or how quickly a load is applied to the disk, is another essential and necessary topic to investigate in future approaches, as high rate impacts can potentially lead to post-traumatic KOA; thus, evaluating the Hippo pathway with various loading rates may provide high valuable insight ^67^.

### The functional interplay of NF-κB and Hippo signaling in chondrocyte mechanoresponses

Several studies have been published investigating the function of the Hippo pathway and YAP/TAZ functionality in articular cartilage homeostasis of knee chondrocytes. Central to the Hippo pathway is the kinase LATS1/2, which functions to phosphorylate YAP/TAZ and inhibit nuclear translocation ^27^. This state of cytoplasmic YAP/TAZ sequestration is referred to as the active state of the Hippo pathway. When the Hippo pathway is inactive, YAP can freely translocate into the nucleus and has been associated with the preservation of articular cartilage integrity specifically via direct interaction with TAK1 and attenuation of the NFκB signaling cascade in an OA mouse model ^27^. The attenuation of NFκB is key to modulating inflammatory pathways associated with OA. In a study by *Yan et* al, using a nanoparticle-based siRNA, NFκB was suppressed and resulted in reduced chondrocyte death and cartilage degeneration ^68^. One study found that YAP is notably activated during the development of OA in mice. The targeted removal of YAP in mouse chondrocytes (conditional knockout or cKO) showed preservation of collagen II expression, a key component of cartilage, thus preventing cartilage degradation in that OA model. Additionally, the introduction of a YAP-specific inhibitor, Verteporfin, through injections directly into the joint, significantly aided in maintaining the balance of cartilage in the OA mouse model ^28^. These apparently conflicting results implied a context-dependent role of YAP and/or Hippo signaling in OA and NFκB *regulation*.

Our results indeed reconciled these studies in the mechanobiology settings. We showed that either full inactivation of Hippo signaling (full YAP activation) or complete inactivation of YAP can block NFκB p65 S536 phosphorylation and target gene expression (**Fig. 5**), suggesting that the full cycle of YAP phosphorylation-dephosphorylation by the Hippo kinase cascade is essential for chondrocyte mechanoresponses, as illustrated in **Fig. 7D**. This notion is further supported by mechanoresponses of PKC and PKA in chondrocytes (**Fig. 7A, Fig. S2A**). Therefore, blocking this Hippo phosphorylation cycle can be a strategy for KOA therapies, as supported by our studies with LATS1/2 inhibitor LATS-IN-1 and PKC inhibitor AEB-071 (**Fig. 6B, Fig. 7B**).

### Characterization of mechanotransduction signaling cascade

Our study has opened a new avenue for identifying and characterizing novel components of mechanotransduction signaling cascade. There are several key mechanism questions to investigate in the next. Firstly, it is worth investigating which PKC isoform(s) responds to mechanical overloading and how mechanical overloading and Hippo signaling regulate the PKC(s). Secondly, there have been two reports on the opposite roles of conventional, novel, and atypical PKCs in Hippo signaling ^69,70^. However, it is unknown how Hippo signaling can in turn regulate PKCs. Our study is therefore the first report showing that Hippo signaling can vice versa regulate PKCs in mechanically loaded cells. Thirdly, conventional, novel, and atypical PKCs can sense various secondary messengers and phosphoinositol ^71^, investigating the functional interactions between PKC and Hippo will likely improve our fundamental understanding of how mechanical loading is sensed at the plasma membranes and subsequently propagated into biochemical signals that eventually alter chondrocyte transcriptomes.

In conclusion, the results presented in the current study suggest that the interplay of NFκB, Hippo, PKC signaling, may be a modifiable target in the response of chondrocytes under mechanical loading, and in a broader context, the response of chondrocytes in patients with KOA. Kinase inhibitors, such as PKC inhibitor AEB-071, which are currently being tested in clinical trials for diseases (i.e., uveal melanoma), can be repurposed for KOA agents that block inflammation and prevent disease progression.

## Materials and Methods

### Preparation of 3-dimensional chondrocyte-agarose constructs

The C28/I2 human chondrocytes were cultured in T-75 flasks with Dulbecco’s Modified Eagle Medium (DMEM, Invitrogen Corp., Carlsbad, CA) supplemented with 10% fetal bovine serum (FBS; Invitrogen Corp.), and 1% antibiotic in an incubator at 37°C and 5% CO_2_. Once cells reached ∼90-95% confluency, they were detached from the flasks using 0.125% trypsin - EDTA solution (Thermo Fisher Scientific Inc, Waltham, MA), centrifuged and resuspended in DMEM to create a cell solution of 2 × 10^7^cells/mL. The cell solution and 4% ultra-low gelling temperature agarose (Sigma, St. Louis, MO) were mixed at a 1:1 ratio to encapsulate 1 × 10^6^ cells in 2% agarose three dimensional (3D) cylindrical constructs (“discs”) (8 mm in diameter, 2 mm in thickness) at 37°C in a hotplate^72^.

### 2D static compression

A stainless-steel cylinder weight (12 mm diameter) of 8.9 g was placed on the top of an agarose disk in contact with chondrocytes cultured on 12 mm Trans-wells with 0.4 μm pore (Fig. 1A) ^35^. In a control group, only the agarose disk was placed on chondrocytes.

### 3D static compression

The 3D discs were each cultured in 2 mL of DMEM supplemented with 10% FBS and 1% antibiotic for 24 hours prior to compressive loading experiments. Samples subjected to compressive loading were placed in a custom-designed bioreactor (**Fig. S1)**. To ensure an equal static compressive strain, a displacement-control approach was used for all testing ^73^.

Strains of 0%, 10%, and 20% were implemented in the dose-response experiment to develop a relationship between strain magnitude and cell mechano-response. At the 20% strain level, we observed increased inflammatory and catabolic gene expression in chondrocytes. Thus, 20% strain was used for the compressive loading experiments to evaluate the Hippo pathway. Furthermore, to perform the compression evaluations the samples were placed in separate custom-made chambers with 600 μL of DMEM. All experiments were conducted in the tissue incubator at 37°C and 5% CO_2_ for a duration of 4 hours. Uncompressed samples were used as control groups.

### Quantitative Real-Time PCR

Immediately following compression, cell-agarose constructs were collected and mechanically lysed in 1 mL of Tri Reagent (Molecular Research Center, Cincinnati, OH) and RNA was extracted per the manufacturer’s instruction. cDNA synthesis was conducted using the Quantabio qScript cDNA synthesis kit (Quanta Biosciences, Gaithersburg, MD). Following cDNA synthesis, quantitative PCR (qPCR) was performed on the samples using the PerfeCta SYBR Green SuperMix (Quanta Biosciences, Gaithersburg, MD). The expression of GAPDH was used as an endogenous control. Gene expression levels were quantified using the 2^−ΔΔCt^ method and normalized to expression levels of respective control group^74^.

**Table 1.**
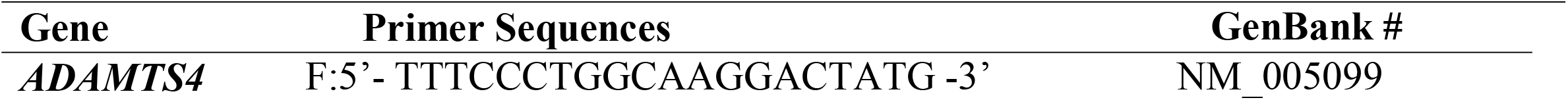
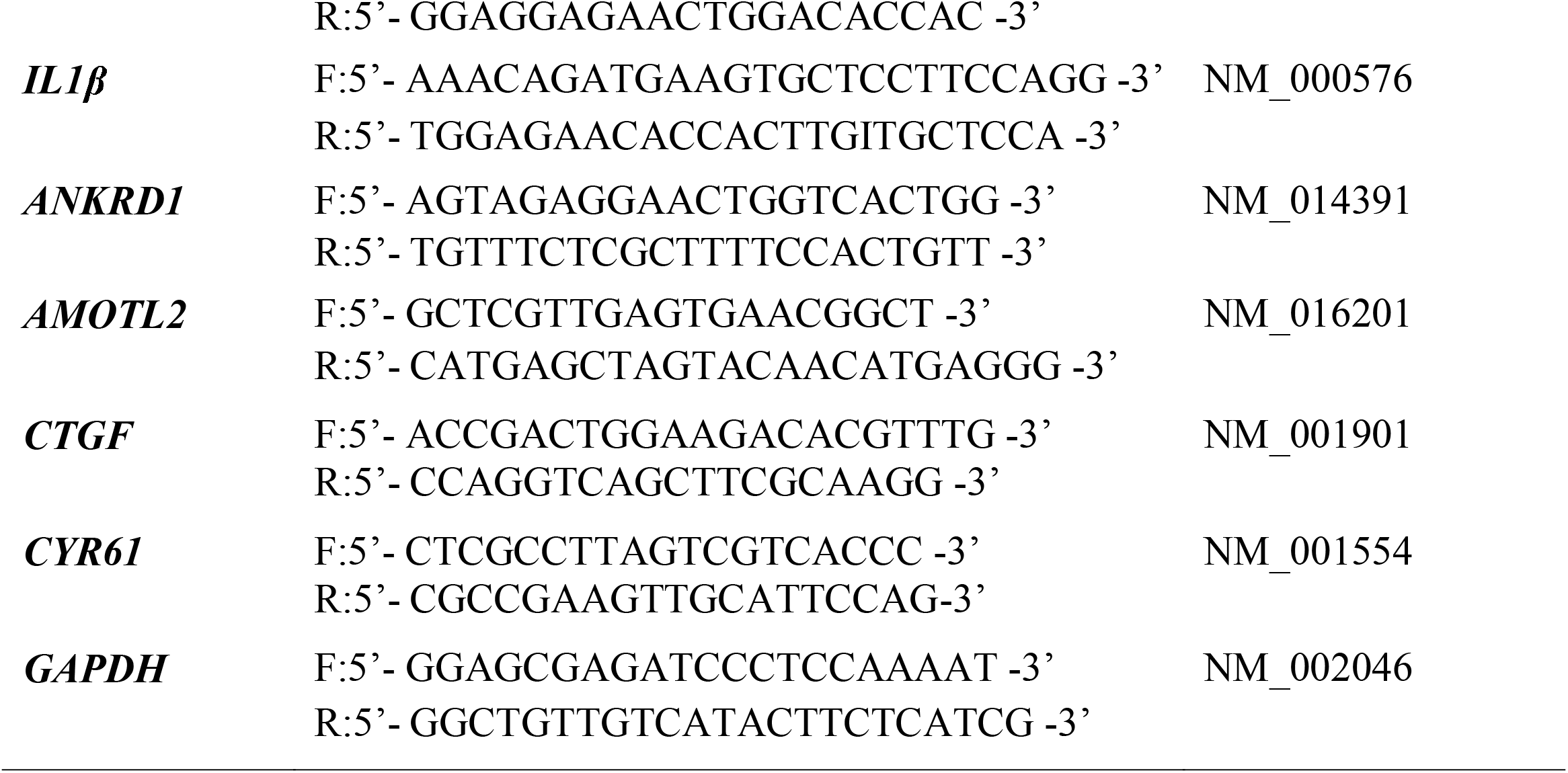
Quantitative real-rime PCR primers.

### CRISPR/Cas9 gene deletion in chondrocytes

LATS1/2 genes were deleted in C28/I2 cells by lentiviral vectors that deliver Cas9 and guide RNAs for LATS1/2 genes into the cells. A LentiCRISPRv2 plasmid with LATS1 guide RNA and a pLenti-Guide-hydro with LATS2 guide RNA were used to package lentiviruses that were later used to infect C28/I2 cells. The lentiviral plasmids were co-transfected in HEK293T cells with PsPAX2 (Addgene#12260) and pMD2.G (Addgene #12259) using PolyJet In Vitro DNA Transfection Reagent (SignaGen, SL100688). Cells were infected for 48 hours, and 2 μg/ml of puromycin and 200 ug/ml of hygromycin were used to select the specific gene knockout cells for 5-6 days, then the cells were maintained in a low concentration of puromycin or hygromycin. The LATS1 and LATS2 gene knockout was confirmed by performing western blot analysis with LATS1 and LATS2 antibodies. Similarly, to evaluate the Hippo pathway activity in the 2D and 3D chondrocyte cultures, YAP or TAZ was also deleted in C28/I2 cells using CRISPR/Cas9.

### Western blot analysis

Either cell-agarose constructs or chondrocytes (monolayer culture) were lysed with 1xSDS PAGE sample buffer. 10 μL protein samples were loaded and separated on 10% SDS PAGE gels, and then were transferred to PVDF membranes and blotted with the antibodies previously described ^54^.

### Statistical analysis

Comparisons between the two groups were done using a student’s two-tailed t-test for the knockout study and 2D *vs* 3D study, while comparisons among three experimental groups were performed using a one-way ANOVA with the Duncan’s Multiple Range Test for the LATS inhibitor study and dose-response study. GraphPad Prism 9 software (GraphPad Software, San Diego, CA) was used for all statistical analyses. In all cases, significance was defined as p ≤ 0.05.

## Funding

C.W., O.F.P., and Q.T.E were supported by the USOAR scholarship from the University of Miami Miller School of Medicine. Z.M. is supported by the National Institute of General Medical Sciences of the National Institutes of Health (NIH) under award number R35GM142504.

## Author contributions

Z.M. and C.-Y. H. conceived the project, obtained the funding for the project, supervised the study, and wrote the manuscript. X.C. generated CRISPR knockout cells, X.C. Y. W., C.W., C. D.R performed biochemical analyses. C.W., O.F.P., L.H., C.F., and Q.T.E. performed the compression experiments. X.C., C.W., O.F.P., C.-Y. H., and Z.M. wrote the manuscript. L.K. and T.M.B. provided intelligent support.

## Competing interests

There is no competing interest from all authors.

---

**Figure S1:**
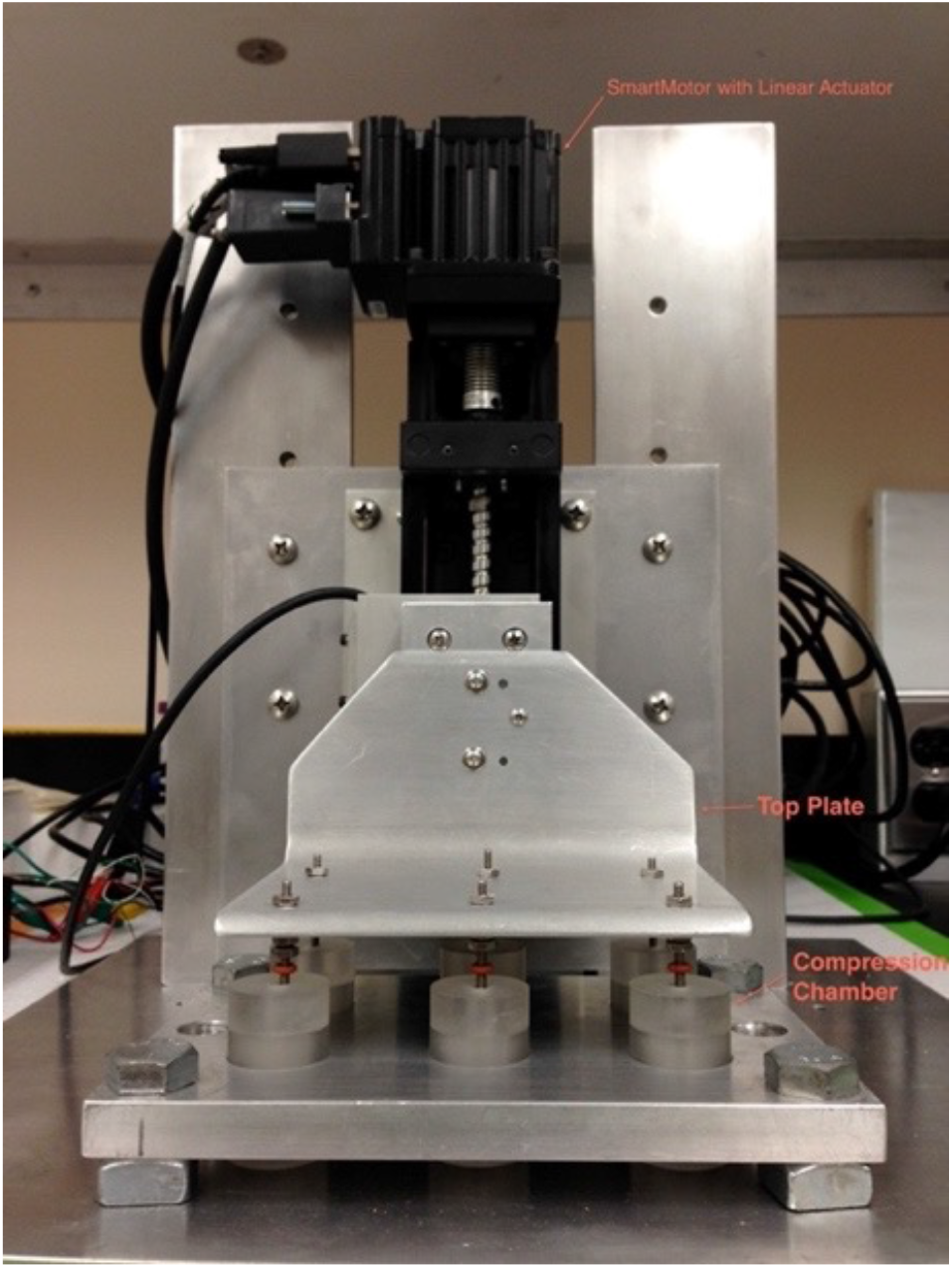
Custom bioreactor for chondrocyte compression. Samples were placed in custom-made chambers. A linear actuator was utilized to apply a static displacement to samples via a top plate to which all compression chambers were connected.

**Figure S2:**
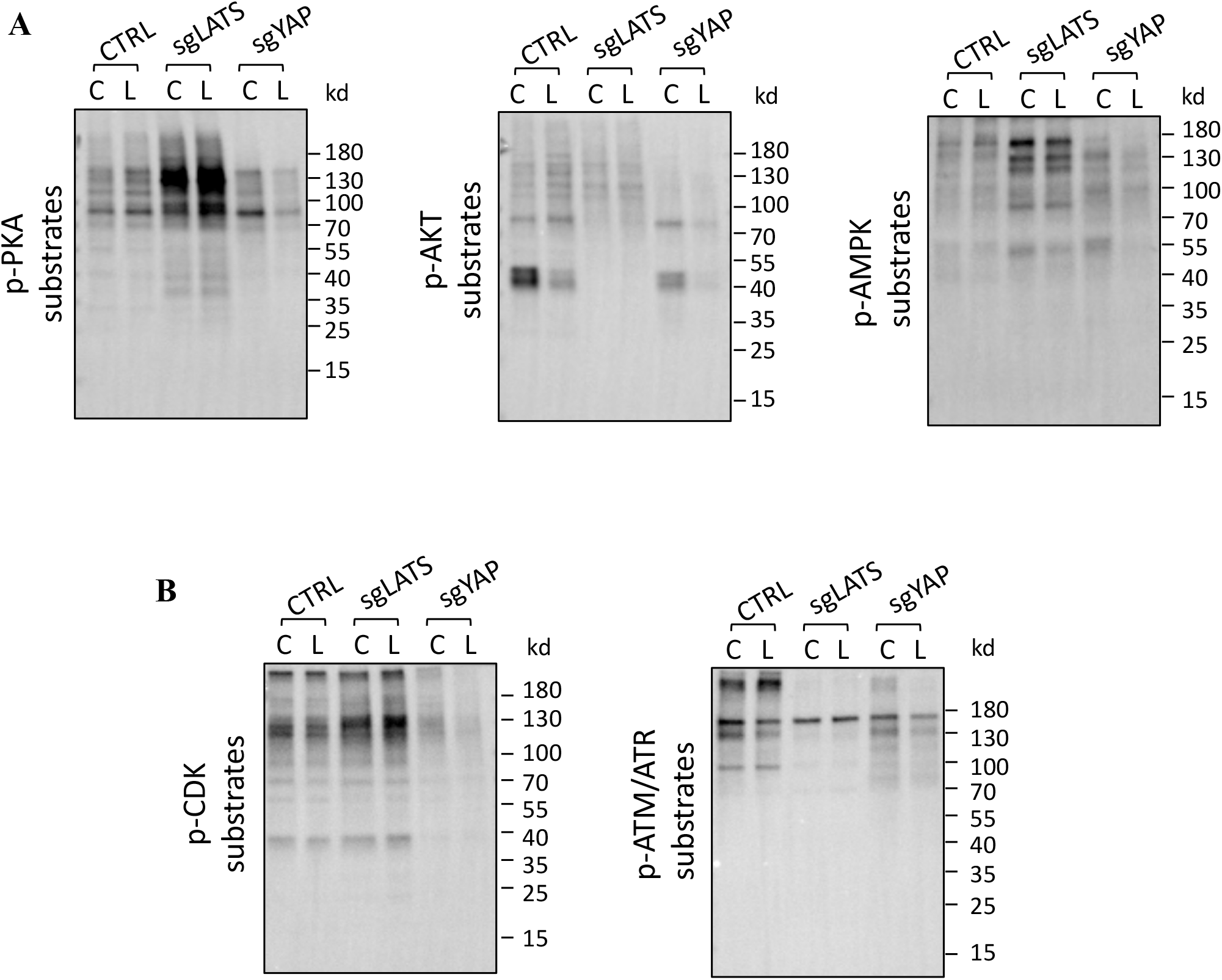
Profiling of activities of protein kinases involved in mechanosensing, metabolism (A), and cell cycle (B), using phosphor suybstrate antibodies for PKA, AKT, AMPK, CDK, and ATM/ATR.

## Notes

### Competing Interest Statement

The authors have declared no competing interest.

## References

1. Martel-Pelletier, J., Barr, A.J., Cicuttini, F.M., Conaghan, P.G., Cooper, C., Goldring, M.B., Goldring, S.R., Jones, G., Teichtahl, A.J., and Pelletier, J.P. (2016). Osteoarthritis. Nat Rev Dis Primers 2, 16072. 10.1038/nrdp.2016.72.

2. Kulkarni, K., Karssiens, T., Kumar, V., and Pandit, H. (2016). Obesity and osteoarthritis. Maturitas 89, 22–28. 10.1016/j.maturitas.2016.04.006.

3. Toivanen, A.T., Heliövaara, M., Impivaara, O., Arokoski, J.P.A., Knekt, P., Lauren, H., and Kröger, H. (2009). Obesity, physically demanding work and traumatic knee injury are major risk factors for knee osteoarthritis—a population-based study with a follow-up of 22 years. Rheumatology 49, 308–314. 10.1093/rheumatology/kep388.

4. Jinks, C., Jordan, K.P., Blagojevic, M., and Croft, P. (2008). Predictors of onset and progression of knee pain in adults living in the community. A prospective study. Rheumatology 47, 368–374. 10.1093/rheumatology/kem374.

5. Tschon, M., Contartese, D., Pagani, S., Borsari, V., and Fini, M. (2021). Gender and Sex Are Key Determinants in Osteoarthritis Not Only Confounding Variables. A Systematic Review of Clinical Data. J Clin Med 10. 10.3390/jcm10143178.

6. King, L.K., March, L., and Anandacoomarasamy, A. (2013). Obesity & osteoarthritis. Indian J Med Res 138, 185–193.

7. Bliddal, H., Leeds, A.R., and Christensen, R. (2014). Osteoarthritis, obesity and weight loss: evidence, hypotheses and horizons - a scoping review. Obes Rev 15, 578–586. 10.1111/obr.12173.

8. Latourte, A., Kloppenburg, M., and Richette, P. (2020). Emerging pharmaceutical therapies for osteoarthritis. Nat Rev Rheumatol 16, 673–688. 10.1038/s41584-020-00518-6.

9. Lee, R., and Kean, W.F. (2012). Obesity and knee osteoarthritis. Inflammopharmacology 20, 53–58. 10.1007/s10787-011-0118-0.

10. Leong, D.J., Hardin, J.A., Cobelli, N.J., and Sun, H.B. (2011). Mechanotransduction and cartilage integrity. Ann N Y Acad Sci 1240, 32–37. 10.1111/j.1749-6632.2011.06301.x.

11. Oliveira, S., Andrade, R., Silva, F.S., Espregueira-Mendes, J., Hinckel, B.B., Leal, A., and Carvalho, O. (2023). Effects and mechanotransduction pathways of therapeutic ultrasound on healthy and osteoarthritic chondrocytes: a systematic review of in vitro studies. Osteoarthritis Cartilage 31, 317–339. 10.1016/j.joca.2022.07.014.

12. Zignego, D.L., Hilmer, J.K., Bothner, B., Schell, W.J., and June, R.K. (2019). Primary human chondrocytes respond to compression with phosphoproteomic signatures that include microtubule activation. J Biomech 97, 109367. 10.1016/j.jbiomech.2019.109367.

13. Zhao, Z., Li, Y., Wang, M., Zhao, S., Zhao, Z., and Fang, J. (2020). Mechanotransduction pathways in the regulation of cartilage chondrocyte homoeostasis. J Cell Mol Med 24, 5408–5419. 10.1111/jcmm.15204.

14. Sophia Fox, A.J., Bedi, A., and Rodeo, S.A. (2009). The basic science of articular cartilage: structure, composition, and function. Sports Health 1, 461–468. 10.1177/1941738109350438.

15. Brown, T.D., Johnston, R.C., Saltzman, C.L., Marsh, J.L., and Buckwalter, J.A. (2006). Posttraumatic osteoarthritis: a first estimate of incidence, prevalence, and burden of disease. J Orthop Trauma 20, 739–744. 10.1097/01.bot.0000246468.80635.ef.

16. Atarod, M., Frank, C.B., and Shrive, N.G. (2015). Increased meniscal loading after anterior cruciate ligament transection in vivo: a longitudinal study in sheep. Knee 22, 11–10.1016/j.knee.2014.10.011.

17. Buckwalter, J.A., Anderson, D.D., Brown, T.D., Tochigi, Y., and Martin, J.A. (2013). The Roles of Mechanical Stresses in the Pathogenesis of Osteoarthritis: Implications for Treatment of Joint Injuries. Cartilage 4, 286–294. 10.1177/1947603513495889.

18. Zhu, J., Zhu, Y., Xiao, W., Hu, Y., and Li, Y. (2020). Instability and excessive mechanical loading mediate subchondral bone changes to induce osteoarthritis. Ann Transl Med 8, 350. 10.21037/atm.2020.02.103.

19. Eckstein, F., Hudelmaier, M., and Putz, R. (2006). The effects of exercise on human articular cartilage. J Anat 208, 491–512. 10.1111/j.1469-7580.2006.00546.x.

20. Sanchez-Adams, J., Leddy, H.A., McNulty, A.L., O’Conor, C.J., and Guilak, F. (2014). The mechanobiology of articular cartilage: bearing the burden of osteoarthritis. Curr Rheumatol Rep 16, 451. 10.1007/s11926-014-0451-6.

21. Akkiraju, H., and Nohe, A. (2015). Role of Chondrocytes in Cartilage Formation, Progression of Osteoarthritis and Cartilage Regeneration. J Dev Biol 3, 177–192. 10.3390/jdb3040177.

22. Kurz, B., Lemke, A.K., Fay, J., Pufe, T., Grodzinsky, A.J., and Schunke, M. (2005). Pathomechanisms of cartilage destruction by mechanical injury. Ann Anat 187, 473–485. 10.1016/j.aanat.2005.07.003.

23. Pan, D. (2010). The hippo signaling pathway in development and cancer. Dev Cell 19, 491–505. 10.1016/j.devcel.2010.09.011.

24. Meng, Z., Qiu, Y., Lin, K.C., Kumar, A., Placone, J.K., Fang, C., Wang, K.C., Lu, S., Pan, M., Hong, A.W., et al. (2018). RAP2 mediates mechanoresponses of the Hippo pathway. Nature 560, 655–660. 10.1038/s41586-018-0444-0.

25. Moya, I.M., and Halder, G. (2019). Hippo-YAP/TAZ signalling in organ regeneration and regenerative medicine. Nat Rev Mol Cell Biol 20, 211–226. 10.1038/s41580-018-0086-y.

26. Dey, A., Varelas, X., and Guan, K.L. (2020). Targeting the Hippo pathway in cancer, fibrosis, wound healing and regenerative medicine. Nat Rev Drug Discov 19, 480–494. 10.1038/s41573-020-0070-z.

27. Deng, Y., Lu, J., Li, W., Wu, A., Zhang, X., Tong, W., Ho, K.K., Qin, L., Song, H., and Mak, K.K. (2018). Reciprocal inhibition of YAP/TAZ and NF-kappaB regulates osteoarthritic cartilage degradation. Nat Commun 9, 4564. 10.1038/s41467-018-07022-2.

28. Zhang, X., Cai, D., Zhou, F., Yu, J., Wu, X., Yu, D., Zou, Y., Hong, Y., Yuan, C., Chen, Y., et al. (2020). Targeting downstream subcellular YAP activity as a function of matrix stiffness with Verteporfin-encapsulated chitosan microsphere attenuates osteoarthritis. Biomaterials 232, 119724. 10.1016/j.biomaterials.2019.119724.

29. Meng, Z., Moroishi, T., and Guan, K.L. (2016). Mechanisms of Hippo pathway regulation. Genes Dev 30, 1–17. 10.1101/gad.274027.115.

30. Cai, X., Wang, K.C., and Meng, Z. (2021). Mechanoregulation of YAP and TAZ in Cellular Homeostasis and Disease Progression. Front Cell Dev Biol 9, 673599. 10.3389/fcell.2021.673599.

31. Fang, T., Zhou, X., Jin, M., Nie, J., and Li, X. (2021). Molecular mechanisms of mechanical load-induced osteoarthritis. International Orthopaedics 45, 1125–1136. 10.1007/s00264-021-04938-1.

32. Meng, Z., Li, F.L., Fang, C., Yeoman, B., Qiu, Y., Wang, Y., Cai, X., Lin, K.C., Yang, D., Luo, M., et al. (2022). The Hippo pathway mediates Semaphorin signaling. Sci Adv 8, eabl9806. 10.1126/sciadv.abl9806.

33. Verma, P., and Dalal, K. (2011). ADAMTS-4 and ADAMTS-5: key enzymes in osteoarthritis. Journal of cellular biochemistry 112, 3507–3514. 10.1002/jcb.23298.

34. Vincent, T.L. (2019). IL-1 in osteoarthritis: time for a critical review of the literature. F1000Res 8. 10.12688/f1000research.18831.1.

35. Tse, J.M., Cheng, G., Tyrrell, J.A., Wilcox-Adelman, S.A., Boucher, Y., Jain, R.K., and Munn, L.L. (2012). Mechanical compression drives cancer cells toward invasive phenotype. Proc Natl Acad Sci U S A 109, 911–916. 10.1073/pnas.1118910109.

36. Fitzgerald, J.B., Jin, M., Dean, D., Wood, D.J., Zheng, M.H., and Grodzinsky, A.J. (2004). Mechanical compression of cartilage explants induces multiple time-dependent gene expression patterns and involves intracellular calcium and cyclic AMP. The Journal of biological chemistry 279, 19502–19511. 10.1074/jbc.M400437200.

37. Rigoglou, S., and Papavassiliou, A.G. (2013). The NF-kappaB signalling pathway in osteoarthritis. Int J Biochem Cell Biol 45, 2580–2584. 10.1016/j.biocel.2013.08.018.

38. Cogswell, J.P., Godlevski, M.M., Wisely, G.B., Clay, W.C., Leesnitzer, L.M., Ways, J.P., and Gray, J.G. (1994). NF-kappa B regulates IL-1 beta transcription through a consensus NF-kappa B binding site and a nonconsensus CRE-like site. J Immunol 153, 712–723.

39. Tian, Y., Yuan, W., Fujita, N., Wang, J., Wang, H., Shapiro, I.M., and Risbud, M.V. (2013). Inflammatory cytokines associated with degenerative disc disease control aggrecanase-1 (ADAMTS-4) expression in nucleus pulposus cells through MAPK and NF-kappaB. Am J Pathol 182, 2310–2321. 10.1016/j.ajpath.2013.02.037.

40. Chang, S.H., Mori, D., Kobayashi, H., Mori, Y., Nakamoto, H., Okada, K., Taniguchi, Y., Sugita, S., Yano, F., Chung, U.I., et al. (2019). Excessive mechanical loading promotes osteoarthritis through the gremlin-1-NF-kappaB pathway. Nat Commun 10, 1442. 10.1038/s41467-019-09491-5.

41. Kumar, A., and Boriek, A.M. (2003). Mechanical stress activates the nuclear factor-kappaB pathway in skeletal muscle fibers: a possible role in Duchenne muscular dystrophy. FASEB J 17, 386–396. 10.1096/fj.02-0542com.

42. Sasaki, C.Y., Barberi, T.J., Ghosh, P., and Longo, D.L. (2005). Phosphorylation of RelA/p65 on serine 536 defines an IkappaBalpha-independent NF-kappaB pathway. The Journal of biological chemistry 280, 34538–34547. 10.1074/jbc.M504943200.

43. Sakurai, H., Chiba, H., Miyoshi, H., Sugita, T., and Toriumi, W. (1999). IkappaB kinases phosphorylate NF-kappaB p65 subunit on serine 536 in the transactivation domain. The Journal of biological chemistry 274, 30353–30356. 10.1074/jbc.274.43.30353.

44. Taniguchi, K., and Karin, M. (2018). NF-kappaB, inflammation, immunity and cancer: coming of age. Nat Rev Immunol 18, 309–324. 10.1038/nri.2017.142.

45. Taylor, R.C., Cullen, S.P., and Martin, S.J. (2008). Apoptosis: controlled demolition at the cellular level. Nat Rev Mol Cell Biol 9, 231–241. 10.1038/nrm2312.

46. Dikic, I., and Elazar, Z. (2018). Mechanism and medical implications of mammalian autophagy. Nat Rev Mol Cell Biol 19, 349–364. 10.1038/s41580-018-0003-4.

47. Vandenabeele, P., Galluzzi, L., Vanden Berghe, T., and Kroemer, G. (2010). Molecular mechanisms of necroptosis: an ordered cellular explosion. Nat Rev Mol Cell Biol 11, 700–714. 10.1038/nrm2970.

48. You, J.S., McNally, R.M., Jacobs, B.L., Privett, R.E., Gundermann, D.M., Lin, K.H., Steinert, N.D., Goodman, C.A., and Hornberger, T.A. (2019). The role of raptor in the mechanical load-induced regulation of mTOR signaling, protein synthesis, and skeletal muscle hypertrophy. FASEB J 33, 4021–4034. 10.1096/fj.201801653RR.

49. Pfister, D., Núñez, N.G., Pinyol, R., Govaere, O., Pinter, M., Szydlowska, M., Gupta, R., Qiu, M., Deczkowska, A., Weiner, A., et al. (2021). NASH limits anti-tumour surveillance in immunotherapy-treated HCC. Nature 592, 450–456. 10.1038/s41586-021-03362-0.

50. Fernando, H.N., Czamanski, J., Yuan, T.Y., Gu, W., Salahadin, A., and Huang, C.Y. (2011). Mechanical loading affects the energy metabolism of intervertebral disc cells. J.Orthop.Res. 29, 1634–1641.

51. Wu, X., Su, J., Wei, J., Jiang, N., and Ge, X. (2021). Recent Advances in Three-Dimensional Stem Cell Culture Systems and Applications. Stem Cells Int 2021, 9477332. 10.1155/2021/9477332.

52. Nukuda, A., Sasaki, C., Ishihara, S., Mizutani, T., Nakamura, K., Ayabe, T., Kawabata, K., and Haga, H. (2015). Stiff substrates increase YAP-signaling-mediated matrix metalloproteinase-7 expression. Oncogenesis 4, e165. 10.1038/oncsis.2015.24.

53. Zhao, B., Li, L., Wang, L., Wang, C.Y., Yu, J., and Guan, K.L. (2012). Cell detachment activates the Hippo pathway via cytoskeleton reorganization to induce anoikis. Genes Dev 26, 54–68. 10.1101/gad.173435.111.

54. Meng, Z., Moroishi, T., Mottier-Pavie, V., Plouffe, S.W., Hansen, C.G., Hong, A.W., Park, H.W., Mo, J.S., Lu, W., Lu, S., et al. (2015). MAP4K family kinases act in parallel to MST1/2 to activate LATS1/2 in the Hippo pathway. Nat Commun 6, 8357. 10.1038/ncomms9357.

55. Dupont, S., Morsut, L., Aragona, M., Enzo, E., Giulitti, S., Cordenonsi, M., Zanconato, F., Le Digabel, J., Forcato, M., Bicciato, S., et al. (2011). Role of YAP/TAZ in mechanotransduction. Nature 474, 179–183. 10.1038/nature10137.

56. Oeckinghaus, A., Hayden, M.S., and Ghosh, S. (2011). Crosstalk in NF-kappaB signaling pathways. Nat Immunol 12, 695–708. 10.1038/ni.2065.

57. Koo, J.H., Plouffe, S.W., Meng, Z., Lee, D.H., Yang, D., Lim, D.S., Wang, C.Y., and Guan, K.L. (2020). Induction of AP-1 by YAP/TAZ contributes to cell proliferation and organ growth. Genes Dev 34, 72–86. 10.1101/gad.331546.119.

58. Abe, J. (2007). Role of PKCs and NF-kappaB activation in myocardial inflammation: enemy or ally? J Mol Cell Cardiol 43, 404–408. 10.1016/j.yjmcc.2007.07.002.

59. Diaz-Meco, M.T., and Moscat, J. (2012). The atypical PKCs in inflammation: NFkappaB and beyond. Immunol Rev 246, 154–167. 10.1111/j.1600-065X.2012.01093.x.

60. Collins, A.T., Kulvaranon, M.L., Cutcliffe, H.C., Utturkar, G.M., Smith, W.A.R., Spritzer, C.E., Guilak, F., and DeFrate, L.E. (2018). Obesity alters the in vivo mechanical response and biochemical properties of cartilage as measured by MRI. Arthritis Res Ther 20, 232. 10.1186/s13075-018-1727-4.

61. Cotofana, S., Eckstein, F., Wirth, W., Souza, R.B., Li, X., Wyman, B., Hellio-Le Graverand, M.P., Link, T., and Majumdar, S. (2011). In vivo measures of cartilage deformation: patterns in healthy and osteoarthritic female knees using 3T MR imaging. Eur Radiol 21, 1127–1135. 10.1007/s00330-011-2057-y.

62. Cantor, J.R., Abu-Remaileh, M., Kanarek, N., Freinkman, E., Gao, X., Louissaint, A., Jr., Lewis, C.A., and Sabatini, D.M. (2017). Physiologic Medium Rewires Cellular Metabolism and Reveals Uric Acid as an Endogenous Inhibitor of UMP Synthase. Cell 169, 258–272 e217. 10.1016/j.cell.2017.03.023.

63. Vernon, L., Abadin, A., Wilensky, D., Huang, C.Y., and Kaplan, L. (2014). Subphysiological compressive loading reduces apoptosis following acute impact injury in a porcine cartilage model. Sports Health 6, 81–88. 10.1177/1941738113504379.

64. Hazbun, L., Martinez, J.A., Best, T.M., Kaplan, L., and Huang, C.-Y. (2021). Antiinflammatory effects of tibial axial loading on knee articular cartilage post traumatic injury. J.Biomech. in press.

65. Li, Y., Frank, E.H., Wang, Y., Chubinskaya, S., Huang, H.H., and Grodzinsky, A.J. (2013). Moderate dynamic compression inhibits pro-catabolic response of cartilage to mechanical injury, tumor necrosis factor-α and interleukin-6, but accentuates degradation above a strain threshold. Osteoarthritis Cartilage 21, 1933–1941. 10.1016/j.joca.2013.08.021.

66. Torzilli, P.A., Bhargava, M., Park, S., and Chen, C.T. (2010). Mechanical load inhibits IL-1 induced matrix degradation in articular cartilage. Osteoarthritis Cartilage 18, 97–105. 10.1016/j.joca.2009.07.012.

67. Argote, P.F., Kaplan, J.T., Poon, A., Xu, X., Cai, L., Emery, N.C., Pierce, D.M., and Neu, C.P. (2019). Chondrocyte viability is lost during high-rate impact loading by transfer of amplified strain, but not stress, to pericellular and cellular regions. Osteoarthritis Cartilage 27, 1822–1830. 10.1016/j.joca.2019.07.018.

68. Yan, H., Duan, X., Pan, H., Holguin, N., Rai, M.F., Akk, A., Springer, L.E., Wickline, S.A., Sandell, L.J., and Pham, C.T. (2016). Suppression of NF-kappaB activity via nanoparticle-based siRNA delivery alters early cartilage responses to injury. Proc Natl Acad Sci U S A 113, E6199–E6208. 10.1073/pnas.1608245113.

69. Gong, R., Hong, A.W., Plouffe, S.W., Zhao, B., Liu, G., Yu, F.X., Xu, Y., and Guan, K.L. (2015). Opposing roles of conventional and novel PKC isoforms in Hippo-YAP pathway regulation. Cell Res 25, 985–988. 10.1038/cr.2015.88.

70. Archibald, A., Al-Masri, M., Liew-Spilger, A., and McCaffrey, L. (2015). Atypical protein kinase C induces cell transformation by disrupting Hippo/Yap signaling. Mol Biol Cell 26, 3578–3595. 10.1091/mbc.E15-05-0265.

71. Parker, P.J., Brown, S.J., Calleja, V., Chakravarty, P., Cobbaut, M., Linch, M., Marshall, J.J.T., Martini, S., McDonald, N.Q., Soliman, T., and Watson, L. (2021). Equivocal, explicit and emergent actions of PKC isoforms in cancer. Nat Rev Cancer 21, 51–63. 10.1038/s41568-020-00310-4.

72. Yin, X., Motorwala, A., Vesvoranan, O., Levene, H.B., Gu, W., and Huang, C.Y. (2020). Effects of Glucose Deprivation on ATP and Proteoglycan Production of Intervertebral Disc Cells under Hypoxia. Sci Rep 10, 8899. 10.1038/s41598-020-65691-w.

73. Fernando, H.N., Czamanski, J., Yuan, T.Y., Gu, W., Salahadin, A., and Huang, C.Y. (2011). Mechanical loading affects the energy metabolism of intervertebral disc cells. J Orthop Res 29, 1634–1641. 10.1002/jor.21430.

74. Genemaras, A.A., Ennis, H., Bradshaw, B., Kaplan, L., and Huang, C.C. (2018). Effects of Anti-Inflammatory Agents on Expression of Early Responsive Inflammatory and Catabolic Genes in Ex Vivo Porcine Model of Acute Knee Cartilage Injury. Cartilage 9, 293–303. 10.1177/1947603516684589.

